# Salivary Proinflammatory Cytokines IL-1β, IL-6 and TNF Decrease With Age

**DOI:** 10.1101/2022.04.25.489450

**Authors:** Evangelina Costantino, Sofía Daiana Castell, María Florencia Harman, María Cristina Pistoresi- Palencia, Adriana Beatriz Actis

**Author notes:** **Author responsible for correspondence** Adriana Beatriz Actis. Instituto de Investigaciones en Ciencias de la Salud (INICSA), CONICET y Facultad de Ciencias Médicas, Universidad Nacional de Córdoba. Blvd. de la Reforma s/n. Ciudad Universitaria. 5000 Córdoba. Argentina., E-mail address., int. 157. E-mail address., int. 157.

## Abstract

**Objective:** to analyze salivary concentrations interleukin-1β (IL-1β), 6 (IL-6) and tumor necrosis factor (TNF) according to age in healthy subjects to determine baseline inflammatory state of the oral mucosa in elderly subjects, who are at higher risk of developing inflammation-related diseases.

**Materials and methods:** sixty-four healthy volunteers were divided into groups according to age: 20-39 (G1; n=20); 40-59 (G2; n=24); 60-80 years (G3; n=20). Their stomatognathic system and periodontal status were assessed; consumption of food sources of fatty acids (FA) was evaluated using a validated food frequency questionnaire. IL-1β, IL-6, and TNF salivary levels were determined (ELISA) in mixed unstimulated saliva. Kruskal Wallis and Spearman’s correlation tests were applied (p<0.05).

**Results:** salivary IL-1β and TNF levels were lower in G2 (p=0.001) and G3 (p<0.001) than in G1; IL-6 levels were lower in G3 than in G2 and G1 (p= <0.001). IL-1β showed the highest concentration in all groups (p<0.001). No statistically significant differences in salivary cytokine levels were observed on sex in any of the groups. Associations were observed between salivary cytokines and consumption of some foods containing FA.

**Conclusion:** salivary concentration of proinflammatory cytokines decreased with age. It could be influenced by ageing-related changes. In addtition, the baseline inflammatory state of the oral mucosa would appear to be influenced by dietary intake of sources of anti-inflammatory FA.

**Clinical relevance:** the knowledge about inflammatory state of the oral mucosa, as determined by salivary cytokine concentrations, could be useful for understanding and prevention of developing inflammation-related diseases.

## INTRODUCTION

Inflammation is an innate immune response that is essential to host homeostasis against invading pathogens and to wound healing [1, 2] and can be acute or chronic. The former is characterized by migration of serum proteins and leukocytes from the blood to extravascular tissues, and recruitment of neutrophils followed by recruitment of monocytes that differentiate into macrophages [2, 3]. These initial events, i.e. vascular permeability and infiltration of leukocytes, are regulated by cytokines and chemokines as well as by lipid mediators deriving from arachidonic acid (AA; 20:4 n-6), like prostaglandins (PGs), leukotrienes (LTs), and endocannabinoids [1, 2], by reactive oxygen species, and by enzyme and amino acid derivatives. Inflammation is usually self-limited and resolves rapidly through negative feedback mechanisms, such as secretion of antiinflammatory cytokines and pro-resolving lipid mediators [2, 4] including resolvins, lipoxins, protectins, and maresins, biosynthesized from n-3 fatty acids (FA) [2, 4, 5].

If timely resolution of inflammation fails, it persists and can progress to chronic inflammation and autoimmune diseases [1, 2]. Interleukins -1β (IL-1β), 6 (IL-6), and tumor necrosis factor (TNF) are important inflammation mediators that initiate and orchestrate a wide range of functional cellular changes, such as induction of inflammation and activation of vascular endothelium, production of PGs, cytokines, and antibodies, recruitment and activation of neutrophils, macrophages, and T and B cells, and promotion of tissue destruction [6, 7].

The oral mucosa acts as a protective barrier and can also be the site of inflammatory processes [8-10]. In the event of physical or chemical damage to the tissue, microbial infection, immune reactions, or malignant transformation [9], keratinocytes and immune cells in the oral mucosa trigger secretion of inflammatory mediators, including cytokines and chemokines, which leads to an inflammatory response [10-12].

Saliva is an important biological fluid that can be used to assess inflammation markers, such as cytokines, secreted in response to activation of the immune system in the oral mucosa [13-15]. It can be collected easily and non-invasively by the patient, and it is useful for diagnosis, monitoring, and follow up of disease and health status [14-16]. Cytokines IL-1β, IL-4, IL-6, IL-8, IL-2, and TNF as well as interferon-γ (IFN-γ), secretory immunoglobulin A (IgAs), α-amylase, cortisol, and total proteins are the most commonly used inflammatory markers [17].

It is also known that ageing affects the homeostatic function of the immune system, causing changes in innate, humoral, and cellular immunity [18, 19], and that it is associated with low grade chronic inflammation caused by mechanical trauma, ischemia, stress, and environmental factors such as ultraviolet radiation. These stimuli induce secretion of molecular patterns associated with pathogens (MPAP), which, via activation of nuclear factor kappa beta (NF-kβ), trigger an inflammatory cascade that leads to an increase in proinflammatory cytokines [20-22]. IL-6, TNF, and IL-1β contribute significantly to unwanted persistence of inflammation, which predisposes to the development of age-related diseases like cancer, rheumatoid arthritis, atherosclerosis, and neurodegenerative diseases [22, 21]. Ageing is characterized by an increase in the concentration of inflammatory mediators in the bloodstream [18, 19, 21], and it is well documented that circulating levels of a number of inflammatory markers and mediators are higher in the elderly than in young subjects. Nevertheless, only a few mediators, namely high-sensitivity reactive protein C, IL-6, IL-10, IL-15, and IL-18 [23], have been shown to be significantly and consistently associated with ageing. Findings regarding other mediators, including IL-1β, serum amyloid protein A, TNF, and IL-8, however, are less consistent [19]. Although the relation between ageing and the immune system, particularly inflammatory processes, has been thoroughly investigated, to our knowledge there are no studies exploring the association between age and inflammation of the oral mucosa as determined by salivary cytokine levels. In view of the above, the aim of the present work was to analyze salivary concentration of proinflammatory cytokines, namely IL-1β, IL-6, and TNF, according to age in healthy men and women in order to determine baseline inflammatory status of the oral mucosa in elderly subjects, who are at higher risk of developing inflammation-related diseases.

## MATERIALS AND METHODS

The study comprised 64 apparently healthy male and female volunteers aged 20 to 80 years recruited among the teaching, technical, administrative, and maintenance staff and students of the School of Dentistry of the National University of Córdoba (UNC). The participants were divided into three groups according to age: 20-39 (G1; n=20), 40-59 (G2; n=24), and 60-80 years (G3; n=20). After obtaining their signed informed consent, clinical records were filled in, and examination of the subjects’ stomatognathic system was performed to determine their general and oral health status. The subjects’ general data as well as those referred to current disease and medication, and the results of their dental-stomatological evaluation were recorded on a standardized form; subjects presenting with infectious disease (visible cavitated carious lesions, periodontal disease, pericoronitis), or systemic or local tumor or metabolic disease, were excluded from the study.

Men and women aged 20 to 80 years with normal body mass index (BMI: 18.5-24.9 kg/m^2^ for subjects aged 20 to 59 years [24] and 23 to 29 kg/m^2^ for subjects aged 60 to 80 years [25], who gave their signed informed consent were included in the study. Smokers, heavy drinkers, subjects on a special diet, who performed intense physical activity, and/or were taking medication (including contraception), subjects presenting with systemic or local infectious, tumoral, or metabolic disease as shown on their clinical chart and by clinical dental-stomatological examination respectively, periodontal disease, and/or exposed to local irritating factors such as removable dentures, dental fracture, and/or an orthodontic appliance, were excluded from the study.

### Examination of the stomatognathic system

The stomatognathic assessment was performed by a single trained dentist and involved examination of both hard and soft tissues of the oral cavity (lips, lip commissures, palate, floor of the mouth, tongue, and cheek mucosa) and salivary glands. The data were recorded on each patient’s clinical record. Periodontal status was also evaluated based on the classification of periodontal disease and conditions established by the American Academy of Periodontology and the European Federation of Periodontology [26, 27]. Examination was performed using a marquis-type probe, and periodontal health was confirmed according to the following parameters: probing depth (PD), clinical attachment level (CAL) ≤ 3mm at all sites, and bleeding on probing<10% [27]. Volunteers presenting with inflammation of the periodontal tissue were excluded.

### Dietary-nutritional data

Given that n-6 and n-3 FA affect the immune system, particularly inflammatory markers [2, 28-30], it was relevant to determine the participants’ eating habits and their dietary intake of sources of FA. To this end, a validated qualitative-quantitative food frequency questionnaire was applied [31]. This structured individual questionnaire provides anthropometric data as well as information on the subject’s dietary pattern. The tool assesses frequency of consumption (monthly, weekly, daily, never) and portion size (small, medium, large) of 257 foods and beverages. Two nutrition specialists administered the questionnaire (blind), which they complemented with a series of photographs to illustrate portion size [32]. The data collected through the questionnaire were processed using *Interfood v*.*1*.*3* software [33], which provides information on food consumption in g/day; we focused on data regarding foods that are a source of FA.

### Saliva collection

Saliva was collected from 8:00 to 10:00 a.m. after a minimum of six to eight hours fasting. With the subject leaning forward, unstimulated whole saliva was collected in sterile plastic test tubes (5 mL or more). Prior to saliva collection, the participants consumed no beverages and remained at rest; they performed no physical activity, and did not brush their teeth or use any oral rinse. The saliva samples were then centrifuged at 3.500 rpm at 4 °C for 15 min, diluted (1:1) in PBS containing protease inhibitors (0.1 mM PMSF, 0.1 mM benzethonium chloride, 10 mM EDTA, 0.01 mg/mL aprotinin A, and 0.05% Tween 20), and frozen at -80 °C until analysis [34].

### Enzyme-linked immunosorbent assay (ELISA)

The following commercially-available kits were used to measure salivary concentration of IL-1β, IL-6, and TNF following the manufacturer’s instructions: set human IL-1β (BD Biosciences, San Diego, USA), set human IL-6 (BD Biosciences, San Diego, USA), and TNF (BD Biosciences, San Diego, USA). The Bradford [35] assay with a BSA standard (Sigma-Aldrich) was used to determine total protein content in saliva. Total protein concentration was used to normalize IL-1β, IL-6 and TNF values for each sample [34]. Salivary levels of cytokines were calculated using *GraphPad Prism 6* software (San Diego, CA, USA) and are expressed as pg/mg.

### Statistical analyses

Salivary levels of the studied cytokines were analyzed according to sex and age, and clinical periodontal parameters, daily food consumption patterns, and dietary intake of sources of FA were analyzed according to age using Kruskal Wallis test. Associations between clinical periodontal parameters and salivary cytokine levels and between dietary intake of sources of FA and salivary cytokine levels were explored using Spearman’s correlation test [36]. A value of p≤0.05 was considered statistically significant. Statistical analysis was performed using Infostat software [37].

## RESULTS

The study population is described in Table 1. Minimum and maximum ages in each group were as follows: G1: 20-34; G2: 41-57; G3: 60-79. No statistically significant differences in total energy value were observed among groups (p>0.05). As shown in Table 2, statistically significant differences in all the clinical periodontal parameters except bleeding on probing were observed among the different age-groups. Of note, given that the total number of teeth was higher in G1 (p<0.001), PD and CAL were determined at a greater number of sites in this group than in G2 and G3. Daily food consumption patterns and consumption of foods that are a source of FA corresponding to each of the studied age-groups are shown in Figure 1 and Table 3. Interestingly, unexpected differences in consumption of fish, seafood, vegetable fat, and legumes were observed between the eldest and the youngest groups (Table 3).

**Table 1.** Study population.

| Characteristics | Age-group |  |  |  |  |  |
| --- | --- | --- | --- | --- | --- | --- |
|  | G1 (n=20) |  | G2 (n=24) |  | G3 (n=20) |  |
| Sex | Women (n=10) | Men (n=10) | Women (n=13) | Men (n=11) | Women (n=10) | Men (n=10) |
| Age (years) | 23.60 ± 2.88 | 26.10 ± 4.18 | 52.00 ± 5.12 | 50.18 ± 5.21 | 67.60 ± 2.55 | 66.90 ± 5.15 |
| BMI | 22.06 ± 1.92 | 23.90 ± 1.17 | 23.45 ± 1.69 | 24.53 ± 0.43 | 27.29 ± 2.08 | 27.45 ± 2.10 |
| TEV (Kcal/day) | 1813.13 ± 313.11 | 2607.47 ± 567.41 | 2055.41 ± 583.56 | 3292.62 ± 1029.92 | 2495.49 ± 890.72 | 2655 ± 622.43 |

The table shows the characteristics of the study population. BMI and TEV data were obtained from the qualitative-quantitative food frequency questionnaire. **G1**: Healthy subjects aged 20-39 years; **G2**: healthy subjects aged 40-59 years; **G3**: healthy subjects aged 60-80 years. **BMI**: Body mass index; **TEV**: total energy value. Values are shown as mean ± standard deviation.

**Table 2.**
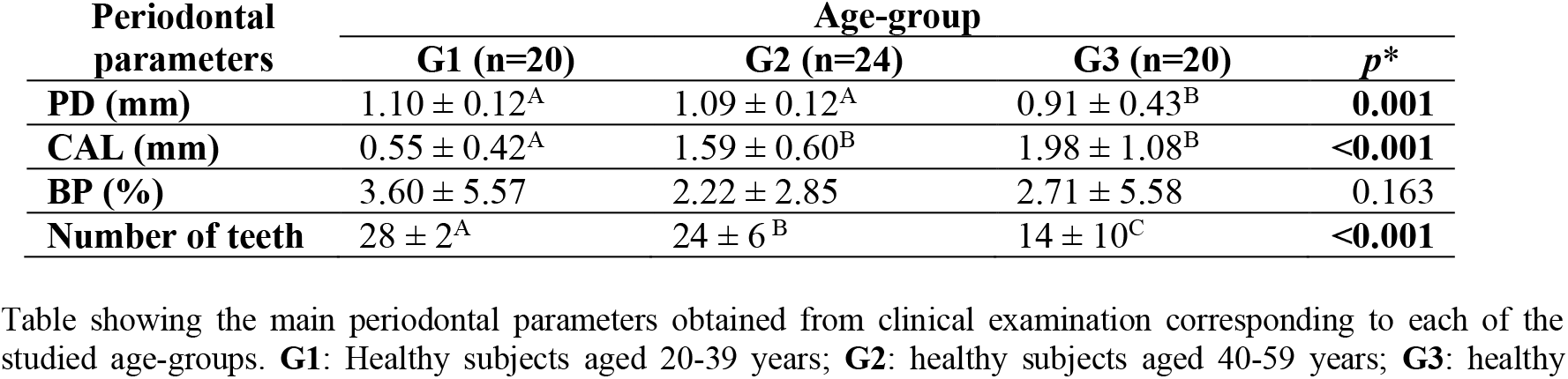
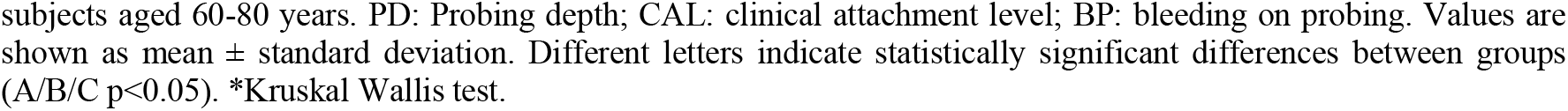
Clinical periodontal parameters corresponding to each age-group.

| Periodontal parameters | Age-group | | | $p^*$ |
| --- | --- | --- | --- | --- |
|  | G1 (n=20) | G2 (n=24) | G3 (n=20) |  |
| PD (mm) | 1.10 ± 0.12<sup>A</sup> | 1.09 ± 0.12<sup>A</sup> | 0.91 ± 0.43<sup>B</sup> | **0.001** |
| CAL (mm) | 0.55 ± 0.42<sup>A</sup> | 1.59 ± 0.60<sup>B</sup> | 1.98 ± 1.08<sup>B</sup> | **<0.001** |
| BP (%) | 3.60 ± 5.57 | 2.22 ± 2.85 | 2.71 ± 5.58 | 0.163 |
| Number of teeth | 28 ± 2<sup>A</sup> | 24 ± 6<sup>B</sup> | 14 ± 10<sup>C</sup> | **<0.001** |

Table showing the main periodontal parameters obtained from clinical examination corresponding to each of the studied age-groups. **G1**: Healthy subjects aged 20-39 years; **G2**: healthy subjects aged 40-59 years; **G3**: healthy
subjects aged 60-80 years. PD: Probing depth; CAL: clinical attachment level; BP: bleeding on probing. Values are shown as mean ± standard deviation. Different letters indicate statistically significant differences between groups (A/B/C $p < 0.05$ ). \*Kruskal Wallis test.

**Table 3.**
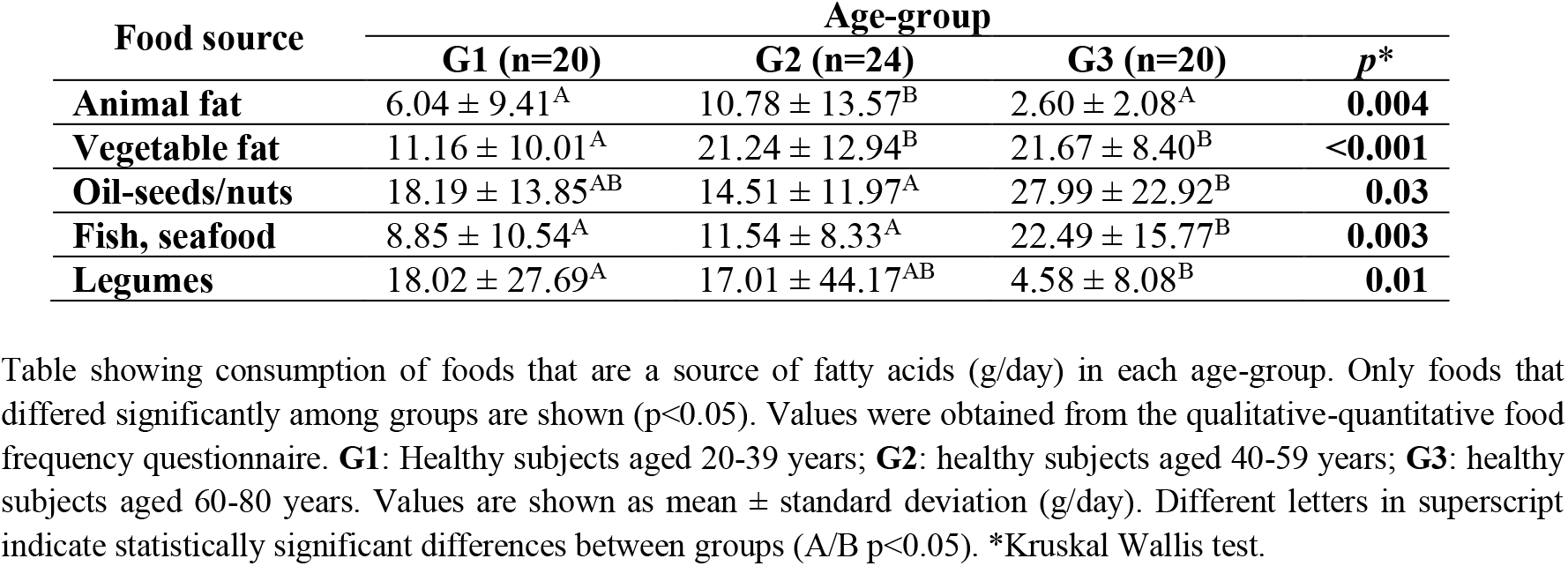
Dietary intake of food sources of fatty acids in each age-group.

Salivary concentration of proinflammatory cytokines IL-1β, IL-6, and TNF according to age and sex are shown in Tables 4 and 5 respectively. It is important to point out that the concentration of these cytokines decreased with age, and were lowest in subjects aged 60 to 80 years (G3). No statistically significant differences in salivary cytokine levels were observed between men and women in any of the three age-groups (p>0.05). Salivary levels of IL-1β were significantly higher than those of IL-6 and TNF (p<0.001) in all three groups; no significant differences were observed between IL-6 and TNF levels.

**Table 4.**
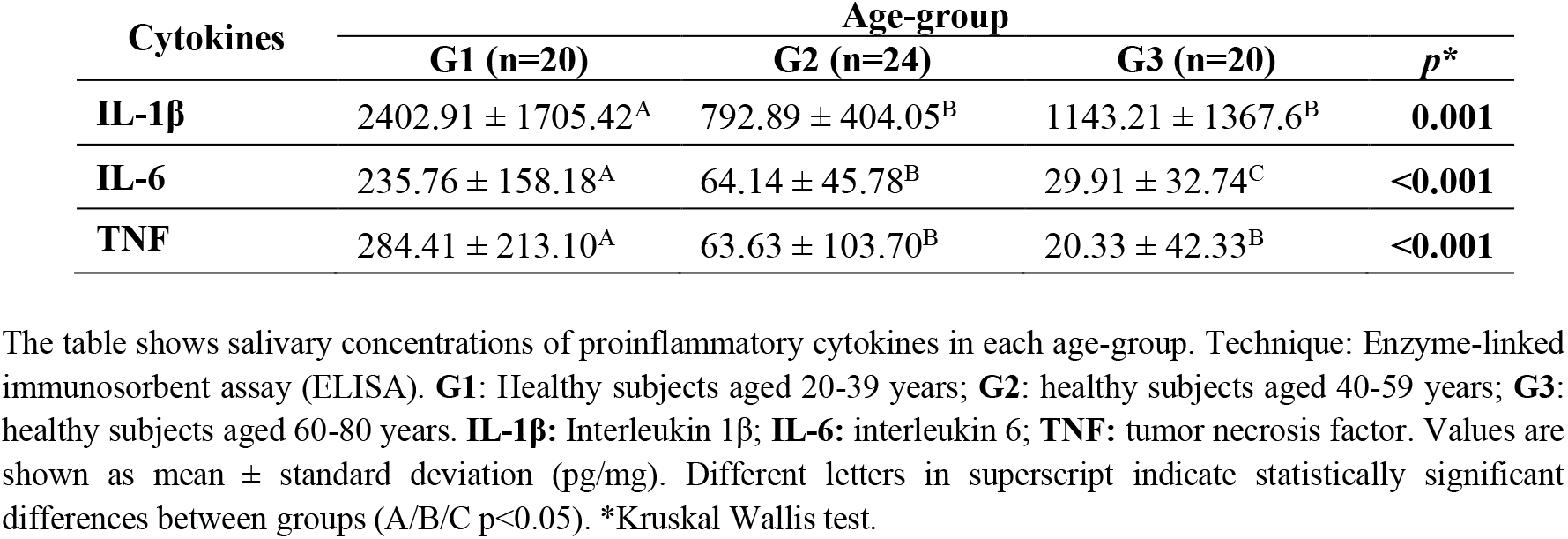
Salivary concentration of cytokines in each age-group.

| Cytokines | Age-group |  |  | *p* * |
| --- | --- | --- | --- | --- |
|  | G1 (n=20) | G2 (n=24) | G3 (n=20) |  |
| **IL-1β** | 2402.91 ± 1705.42<sup>A</sup> | 792.89 ± 404.05<sup>B</sup> | 1143.21 ± 1367.6<sup>B</sup> | **0.001** |
| **IL-6** | 235.76 ± 158.18<sup>A</sup> | 64.14 ± 45.78<sup>B</sup> | 29.91 ± 32.74<sup>C</sup> | **<0.001** |
| **TNF** | 284.41 ± 213.10<sup>A</sup> | 63.63 ± 103.70<sup>B</sup> | 20.33 ± 42.33<sup>B</sup> | **<0.001** |

The table shows salivary concentrations of proinflammatory cytokines in each age-group. Technique: Enzyme-linked immunosorbent assay (ELISA). **G1**: Healthy subjects aged 20-39 years; **G2**: healthy subjects aged 40-59 years; **G3**: healthy subjects aged 60-80 years. **IL-1 $\beta$** : Interleukin 1 $\beta$ ; **IL-6**: interleukin 6; **TNF**: tumor necrosis factor. Values are shown as mean ± standard deviation (pg/mg). Different letters in superscript indicate statistically significant differences between groups (A/B/C $p < 0.05$ ). \*Kruskal Wallis test.

**Table 5.**
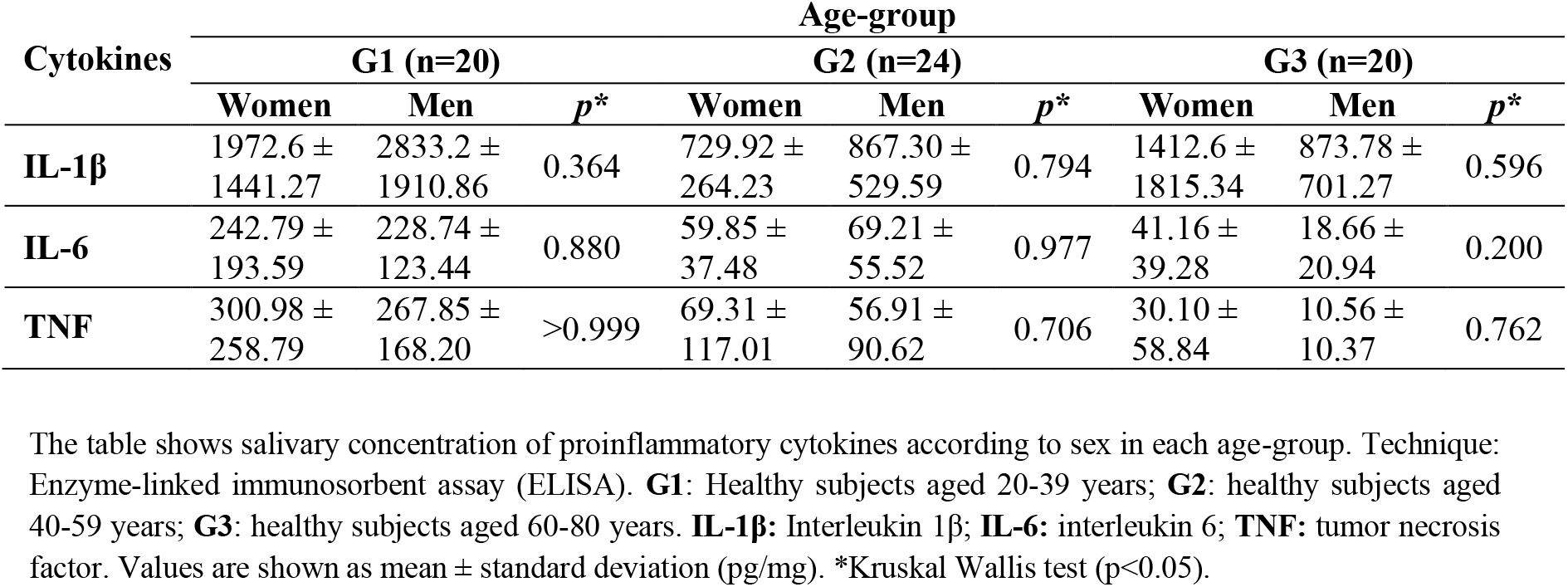
Salivary concentration of proinflammatory cytokines according to sex.

No association was observed between salivary concentration of the proinflammatory cytokines and the clinical periodontal parameters studied here (p>0.05).

Interestingly, associations were observed between consumption of some foods that are a source of FA, namely fish, seafood, pork, beef, entrails, animal fat, vegetable fat, legumes, whole-fat dairy products, and low-fat dairy products, and each of the studied age-groups (Table 6). Only correlations showing statistically significant differences among the studied age-groups were calculated (Table 3). No association between cytokines and BMI as well as TEV was observed in any group (p>0.05).

**Table 6.**
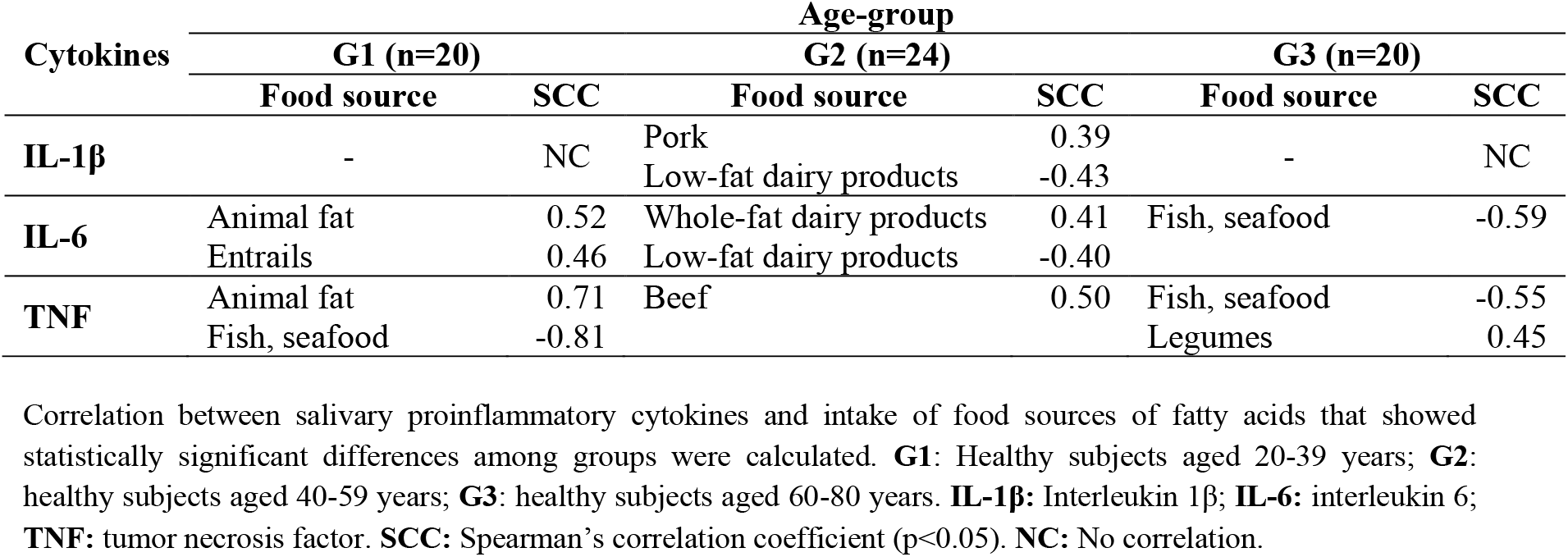
Correlation between salivary proinflammatory cytokines and consumption of food sources of fatty acids.

| Cytokines | Age-group |  |  |  |  |  |
| --- | --- | --- | --- | --- | --- | --- |
|  | G1 (n=20) |  | G2 (n=24) |  | G3 (n=20) |  |
|  | Food source | SCC | Food source | SCC | Food source | SCC |
| **IL-1β** | - | NC | Pork | 0.39 | - | NC |
|  |  |  | Low-fat dairy products | -0.43 |  |  |
| **IL-6** | Animal fat | 0.52 | Whole-fat dairy products | 0.41 | Fish, seafood | -0.59 |
|  | Entrails | 0.46 | Low-fat dairy products | -0.40 |  |  |
| **TNF** | Animal fat | 0.71 | Beef | 0.50 | Fish, seafood | -0.55 |
|  | Fish, seafood | -0.81 |  |  | Legumes | 0.45 |

Correlation between salivary proinflammatory cytokines and intake of food sources of fatty acids that showed statistically significant differences among groups were calculated. **G1**: Healthy subjects aged 20-39 years; **G2**: healthy subjects aged 40-59 years; **G3**: healthy subjects aged 60-80 years. **IL-1 $\beta$** : Interleukin 1 $\beta$ ; **IL-6**: interleukin 6; **TNF**: tumor necrosis factor. **SCC**: Spearman's correlation coefficient ( $p < 0.05$ ). **NC**: No correlation.

## DISCUSSION

To our knowledge, this is the first study to analyze salivary concentration of proinflammatory cytokines IL-1β, IL-6, and TNF of healthy subjects according to age and sex in order to know about the baseline inflammatory state of the oral mucosa of elderly subjects, who are at higher risk of developing inflammation-related diseases. We found the eldest subjects to show lower concentrations of IL-1β, IL-6, and TNF as compared to the younger age-groups. Our results would suggest that ageing may be associated with a decrease in inflammation of the oral mucosa as determined by salivary concentration of proinflammatory cytokines. Of note, there is limited research on the relation between ageing and salivary concentration of cytokines.

Bäck et al. found positive correlation between age and salivary levels of inflammation marker PGE2, and higher salivary MMP-9 activity in subjects over the age of 40 years, irrespective of sex [38]. As regards the latter factor, our results showed no differences in salivary concentrations of IL-1β, IL-6, or TNF between male and female participants, which would suggest that there is no association between sex and the concentration of the inflammation markers studied here. Riis et al. analyzed cytokines IFNg, TNF, IL-1β, IL-2, IL-6, IL-8, IL-10, and IL-12p70 in serum and saliva in adolescent girls aged 13 to 17 years across three consecutive annual assessments. They observed a negative association between age and salivary levels of most of the studied cytokines (older subjects showed lower cytokine levels), suggesting a relation between inflammatory state of the oral mucosa and hormonal changes associated with pubertal development in adolescent girls. As in our study showing higher salivary concentrations of IL-1β than of IL-6 and TNF, the authors found salivary IL-1β and IL-8 levels to be 20-fold higher compared to the remaining studied cytokines [39]; it must be pointed out, however, that the studies are not comparable due to the different ages of the study populations. Of note, unlike our study, the methodology used in the reports mentioned above did not take into account the presence of periodontal disease, exposure to local irritants such as removable dentures and tooth fracture, dietary intake of sources of FA with anti-inflammatory and proinflammatory properties, special food diets, heavy alcohol drinking, medication, or other factors known to affect inflammation.

There are numerous studies on salivary cytokines [40-43]; nevertheless, none were conducted in systemically and stomatognathically healthy patients, and none took age and sex into account.

Saliva contains water, proteins, bioactive peptides, nucleic acids, lipids, and electrolytes, in addition to complementary contributions from the oral mucosa, periodontium, oral microbiota and its metabolites, blood cells, and exfoliated epithelial cells [14, 44, 45]. Most components of saliva are produced locally by the salivary glands, but some molecules pass from the blood to saliva through the salivary ducts and via biological processes such as passive diffusion, active transport, and ultrafiltration [44, 45]. It is noteworthy, however, that the concentration of most of the serum components of saliva is approximately 300 to 3000 times lower in saliva than in serum.

A number of studies have described salivary gland production of proteins. Amino acids enter the acinar cells by active transport, and after synthesis, most intracellular proteins are stored in granules that are released upon secretory stimulation [16, 46, 47]. However, there are no studies analyzing salivary gland production of cytokines nor on the proteins derived from the local vasculature. It is therefore relevant to further investigate the contribution of the salivary glands and of the local vasculature to the final concentration of these cytokines in saliva.

The decrease in salivary concentrations of proinflammatory cytokines in elderly subjects may be due to different age-related changes, such as a reduction in microvasculature, which would lead to a decrease in components of saliva derived from the blood resulting from a diminution in passive diffusion, ultrafiltration, and active transport; a decrease in the total concentration of salivary proteins [48]; and anatomical and histological changes in salivary glands, such as acinar cell atrophy and replacement of normal parenchyma with fibrosis [48, 49]. According to Xu et al., the salivary glands of 23-year-old subjects have a lobular structure and a more uniform and compact parenchyma as compared to the salivary glands of 83-year-olds, who show a proportional increase in adipose and fibrous tissue and a decrease in the volume of acinar cell secretion, which could cause salivary gland hypofunction. In addition to these structural and functional changes, the intensity of stimulation and reflex decreases with age as a result of a decrease in olfactory and taste sensitivity and diminished neuronal saliva stimulation and blood perfusion in the salivary glands [49]. Although it is well documented that the composition of saliva changes in subjects over the age of 60 years -increase in potassium, chloride, phosphate, uric acid, lysozyme, amylase, and IgAs, decrease in calcium, lactoferrin, transferrin, oxidized glutathione, and decreased peroxidase activity-[49], little is known about the relation between salivary concentrations of cytokines and age.

Ageing is associated with low grade chronic inflammation [20] and increased serum concentration of proinflammatory molecules [18, 19, 21]. Although numerous studies have shown that circulating levels of a number of inflammation mediators are higher in the elderly than in young individuals [19, 21], only some serum mediators have been shown to be significantly and consistently associated with age [19]. Few studies have analyzed age-related changes in inflammation mediators in saliva, and results are not conclusive [39]. In this regard, Riis et al. found no significant correlation between salivary and serum concentrations of cytokines IFNg, TNF, IL-2, IL-6, IL-8, IL-10, and IL-12p70 in adolescent girls aged 13 to 17 years. The authors also reported lower levels of all the studied cytokines, except for IL-8 and IL-1β, in saliva than in serum and a positive association between salivary and serum IL-1β levels [39]. Nam et al. showed that salivary levels of IL-1β and IL-6 were higher than TNF levels in young men, and only found salivary IL-6 to correlate significantly, albeit weakly, with its serum homologue [43].

Reports in the literature and the findings shown here suggest that although ageing is associated with low grade chronic inflammation, as evidenced by the increase in serum inflammatory markers, as are cytokines [20, 21], the same would not seem to apply to salivary concentrations of inflammation markers, which could be related with the aforementioned ageing-related histological and functional changes in the salivary glands.

In order to protect the underlying tissues from different noxae, the oral mucosa produces cytokines and chemokines (via keratinocytes) that promote cellular infiltration, vascular response, tissue destruction, and cell proliferation [10-12]. This protective mucosa is endowed with cells from the innate immune system cells, as are macrophages, dendritic cells, Langerhans cells, natural killer cells, polymorphonuclear leukocytes and their associated inflammatory mediators, including cytokines, chemokines, antibacterial peptides, and components of the complement system. IgAs, oral keratinocyte-derived biologic mediators, and components of the gingival crevicular fluid contribute an additional dimension to oral mucosal immunity [8, 10, 12]. Thus, the mucosa transudate would supply the saliva with cytokines through activation of the immune system of the oral cavity. Ageing-related changes include thinning of the mucosal epithelium together with thickening of the keratin layer and decreased microvasculature [48], all of which could interfere in the production and passage of cytokines into the oral cavity in the elderly, leading to a decrease in the concentration of salivary proteins.

Changes in salivary concentrations of proinflammatory cytokines may be related with consumption of food sources of FA. Polyunsaturated FA (PUFA) of the n-6 series have proinflammatory effects, whereas those of the n-3 series, like docosahexaenoic acid (DHA) and eicosapentaenoic acid (EPA), have immunomodulation and antiinflammatory effects [2, 28-30] by down-regulating NF-kβ, IL-1β, TNF and IL-6 and up-regulating IL-10 [19, 50], and subsequently altering proinflammatory cytokine transcription. In addition, there are bioactive compounds derived from EPA and DHA, including resolvins, protectins, maresins, and PGs E3, which are antiinflammatory lipid mediators that exert their effect through different mechanisms of action [28, 29]. An increase in n-3 FA alters the fatty acid composition of the membrane phospholipids of immune cells, affecting the proinflammatory signaling pathway [29]. In view of this association, we analyzed consumption of foods that are a source of FA as a variable potentially affecting salivary concentration of cytokines.

Participants in the eldest age-group reported higher consumption of fish, seafood (rich in DHA and EPA), and vegetable fat (rich in n-3 and n-6 monounsaturated fatty acids-MUFA and PUFA) than participants in the youngest age-group, and lower consumption of animal fat (rich in saturated FA and n-6 PUFA) as compared to participants in the middle age-group. Our results showed a negative association between consumption of fish and seafood and IL-6 and TNF concentration in saliva in the eldest and youngest age-groups, whereas consumption of animal fat correlated positively with salivary IL-6 and TNF in the youngest age-group.

Although the associations between consumption of foods containing DHA and EPA and salivary levels of the studied cytokines shown in our study are interesting from a nutritional and immunological point of view, daily consumption of these foods was relatively low in the eldest age-group. It is therefore relevant to further investigate this association in individuals who have a higher consumption of foods containing FA with antiinflammatory effects, than that self-reported by the participants of this study.

To conclude, the present study analyzing saliva samples of healthy women and men showed that salivary levels of proinflammatory cytokines IL-1β, IL6, and TNF decrease with age. According to this finding, the salivary concentration of cytokines could be influenced by ageing-related changes, such as morphological and functional changes in the salivary glands, structural changes in the mucosal epithelium, and decreased microvasculature. In addition, the baseline inflammatory state of the oral mucosa would appear to be influenced by dietary intake of foods that are a source of FA with antiinflammatory effects. Further studies are necessary to confirm the findings reported.

## DECLARATIONS

### AUTHOR CONTRIBUTIONS

All authors contributed to the study conception and design. Data collection was carried out by Evangelina Costantino. Material preparation and analysis were performed by Evangelina Costantino, Sofía Daiana Castell, María Florencia Harman, María Cristina Pistoresi-Palencia and Adriana Beatriz Actis. The original manuscript was written by Evangelina Costantino and all authors worked on it in reviewing and editing as well as they read and approved the final version.

### Ethical approval

All procedures performed in studies involving human participants were in accordance with the ethical standards of the institutional and/or national research committee and with the 1964 Helsinki declaration and its later amendments or comparable ethical standards. The study was approved by the Institutional Committee of Ethics in Health Research (CIEIS) of the School of Dentistry, National University of Cordoba, Argentina (ODO-CIEIS N°6T/15).

### Informed consent

Written informed consent was obtained from all subjects included in this study.

## ACKNOWLEDGMENT

Authors wish to acknowledge the assistance of the Consejo Nacional de Investigaciones Científicas y Tecnológicas (CONICET) (Res 574/19 APN-DIR#CONICET) and the Secretaría de Ciencia y Tecnología of the Universidad Nacional de Córdoba (Res N° 411/18), for funding research.

